# Synergistic role of riboflavin-auxotrophic *Enterococcus* for MR1 expression and intra-tumoral mucosal associated invariant T (MAIT) cell activation

**DOI:** 10.1101/2025.11.25.690539

**Authors:** Pakhi Birla, Lansaol Yang, Wanting Shan, Sakura Minamisawa, Omkar Dhaygude, Haritha Manoj, Ying Zheng, Hongni Fan, Hadley Beauregard, Kellie N. Smith, Fyza Y. Shaikh, Cynthia L. Sears, Drew M. Pardoll, Franck Housseau

## Abstract

The role of mucosal invariant T cells (MAITs), in the lung tumor microenvironment remains poorly understood, especially in the setting of immune checkpoint inhibitors. We identified intratumoral MAIT cells from paired single cell RNA and TCR sequencing datasets of tumor infiltrating CD3 T cells isolated from non-small cell lung cancer tumors in patients receiving neoadjuvant PD-1 blockade therapy. MAIT cells were subclustered to identify conventional MAIT-associated TCR clonotypes predicted to recognize intratumoral bacteria, which we then tested for functional recognition using a MAIT TCR capture functional assay. Strikingly, although not directly recognized by MAIT cells, previously identified probiotic *Enterococcus spp* and detected in the intratumoral microbiome of lung cancer patients, selectively synergized with exogenous riboflavin biosynthesis-derived metabolites to induce expression of MR1 by antigen presenting cells, including dendritic cells, B cells and mononuclear phagocytes. Boosting of MR1 cell surface expression resulted from perturbation of endo-lysosomal vacuolar pathway by *Enterococcus* and recycling of early endosomal MR1 to the cytoplasmic membrane. Riboflavin auxotrophic *Enterococcus spp* may therefore exercise their beneficial immunomodulatory functions upon immune checkpoint blockade treatment, at least in part, by promoting intratumoral MR1 expression and innate like T cell activation. Our results indicate that composition of the intratumoral microbiome during immune checkpoint inhibitor treatment has the potential to impact the function of human intratumoral MAIT cells.

## INTRODUCTION

The mucosal associated invariant T (MAIT) cell subset is a frequent innate immune population residing in mucosal sites, liver, and blood (Wong et al., 2017). MAIT cells can rapidly react to disruption of epithelium, mechanical and infectious, to maintain mucosal homeostasis. This early response positions MAIT cells as a critical first line of defense against aggressions (Constantinides et al., 2019). MAIT cells are characterized by functional versatility allowing them to recognize and kill infected epithelial cells (Le Bourhis et al., 2013; Le Bourhis et al., 2010) while also contributing to wound healing and tissue repair response (Bugaut et al., 2024; Germain et al., 2025). They are characterized by semi-invariant TCR alpha chains with a canonical gene recombination TCRAV1-2 / J33, J12 or J20, and paired with restricted set of beta chains (predominantly TCRB V6 or 20) (Reantragoon et al., 2013), MAIT cells are part of a broader group of highly heterogeneous innate-like T cells which recognize target cells through the interaction of their TCR with the MHC 1b molecules, MHC-class-I-related protein 1 (MR1) (Koay et al., 2019). Their development in early-life and their functions at the epithelial barriers are mostly related to the regulation of MR1 expression by bacterial ligands. Specifically, 5-(2-oxopropylideneamino)-6-D-ribitylaminouracil (5-OP-RU) and 5-(2-oxoethylideneamino)-6-D-ribitylaminouracil (5-OE-RU) derived from non-enzymatic reactions between bacterial riboflavin biosynthesis pathway intermediates 5-amino-6-D-ribitylaminouracil (5-A-RU) and methylglyoxal or glyoxal, respectively (Constantinides et al., 2019; Corbett et al., 2014). The crosstalk between MAIT cells and bacteria is therefore an important contributor to the fitness of mucosal immunosurveillance (Constantinides et al., 2019).

Non-MAIT, MR1-restricted T cells recognize a variety of unrelated host-derived MR1 ligands associated with different cell types, tissues and pathological contexts such as infection, cancer and metabolism. (Crowther et al., 2020; Dolton et al., 2025; Gherardin et al., 2016; Ito et al., 2024; Keller et al., 2017; Lepore et al., 2017; Meermeier et al., 2016). Such ligands include cholic acid 7 sulfate (CA7S), derived from bile acid metabolism (Ito et al., 2024), or tumor-associated nucleobase-derived compounds (e.g. carbonyl adduct of adenine called M3Ade), which both can trigger polyclonal MR1-restricted T cell activation (Dominguez et al., 1985; Vacchini et al., 2024). Strikingly, healthy individuals and cancer patients possess MR1-restricted T cells which recognize cancer cells but not 5-OP-RU (Dolton et al., 2025).

Analysis of single cell transcriptomics and TCR sequencing of tumor-infiltrating lymphocytes (TIL) generated from patients with lung cancer receiving neoadjuvant anti-PD1 showed that MAIT cells represented a sizeable group of innate cells (Caushi et al., 2021). In one patient with complete response, MAIT cells exhibited the strongest clonal expansion. Although MAIT cell functions can be altered by PD-1 signaling and blockade of this immune checkpoint can restore their cytotoxic activity in vitro (Ruf et al., 2023), it is unclear whether MAIT cells can contribute to anti-tumor responses in the setting of immune checkpoint blockade (ICB).

One outstanding questions is whether the remarkable variability of abundance of MAIT cells across individuals, a characteristic attributed, at least in part, to the interindividual diversity of commensal flora (Constantinides et al., 2019), may be contributing to the differential sensitivity of patients to ICB (Qu et al., 2024; Shi et al., 2023). Herein, we analyzed the crosstalk between bacteria and MAIT cells to understand how lung intratumoral bacteria can modulate the activation of clonally expanded MAIT cells. Our results showed that riboflavin-auxotrophic *Enterococcus* species (*Enterococcus spp*) demonstrated a unique capacity in boosting intratumoral MAIT cell activation by 5-OP-RU via the selective upregulation of MR1 expression on antigen presenting cells (APC, e.g. B cells, dendritic cells and monocytic cell lines). Importantly, this synergistic activity differed by species. *E. hirea*, *E. faecalis* and *E. faecium* activated MAIT cells in combination with 5-OP-RU, while *E. avium* did not, suggesting that species-level genome coded factors, rather than pathogen-associated membrane patterns, are contributing to the synergistic upregulation of MR1 expression on APCs. We found that the synergistic upregulation of MR1 was not transcriptionally programmed but rather was dependent on the alteration of the endo-lysosomal intracellular compartment by *Enterococcus spp* and increased MR1 recycling at the surface of APCs.

The discovery that specific components of tumor microbiome can synergize with vitamin-precursor metabolites to up-regulate MR1 expression on APCs and boost local innate immunity is unveiling new opportunities for modulating immune surveillance and improving response to cancer immunotherapy, especially for ICB.

## RESULTS AND DISCUSSION

### Intratumoral clonal expansion of conventional MAIT (_conv_MAIT) in patients with lung cancer treated with neoadjuvant PD-1 blockade

To identify innate immune cell populations in the lung tumor microenvironment (TME), we analyzed previously published single cell transcriptomic datasets from patient-matched tumor and distant normal lung tissues (Caushi et al., 2021). The initial CD3+ T cell clustering identified a MAIT cell cluster representing **6.3**% of all T cells (9% in both Normal (n=12) and Tumor (n=15) among other tissues) collected from 16 patients (7 pathological responses [R] and 9 without responses [NR]) (Caushi et al., 2021). MAIT cells represented 8.6% of tumor-infiltrating T cells in R patients and 9.6% in NR patients. MAIT-annotated cluster was further sub-clustered into **CD4^neg^CD8^neg^ cytotoxic MAIT1** (*PRF1^pos^, IFNGR1^pos^*), **CD4^pos^ Naïve MAIT1** (*CD62L^pos^, CCR7^pos^, SOCS1^pos^*), **CD4^pos^ MAIT17** (*MAF^pos^, CCR6^pos^, BATF^pos^, RORA^pos^, AHR^pos^, PTPN13^pos^*), **CD8^pos^ activated MAIT1** (*HLA-DRB1^pos^, PRF1^pos^, GZMA^pos^, GZMB^pos^,*), **tissue resident memory** (**TRM) MAIT1** (CD69*^pos^*, CD62L*^pos^*, IL7R*^pos^*, KLRG1*^pos^*, TXNIP*^pos^*, ADBR2*^pos^*, ARRDC3*^pos^*, STAT1*^pos^*, SOCS1*^pos^*, IFNGR1*^pos^*) and **activated CD8*^pos^* TRM MAIT cells** (*HLA-DRB1^pos^*, ITGAE*^pos^*, LINC02446*^pos^*, *SPRY1^pos^, ZEB2^pos^, PRF1^pos^, GZMA^pos^, GZMB^pos^*) (**Fig.1A and B**). Noteworthy, we observed different densities of cells within the *cytotoxic MAIT1* cluster between R and NR, as well as a higher density of cells in the *MAIT17* and *Activated tissue resident MAIT* clusters in R patients (**Fig.1B**). Furthermore, while c*ytotoxic MAIT1* cluster in the tumor of R patients had a significantly higher expression of cytotoxicity associated genes such as PRF1, NCR3 and NKG7 (**Fig.S1A**), the same MAIT cluster in NR patients was characterized by higher expression of the immunosuppressive TGFB activation pathway (*TGFB* and *ITGVA*)(Malenica et al., 2021). These results suggest that the transcriptional profile of MAIT cells could be reflective of the patient’s response to ICB. We herein paid attention to how the intratumoral bacteria and their metabolites, biosynthetic precursors of Riboflavin (VitB2), can impact _conv_MAIT cells activation.

**Figure 1.**
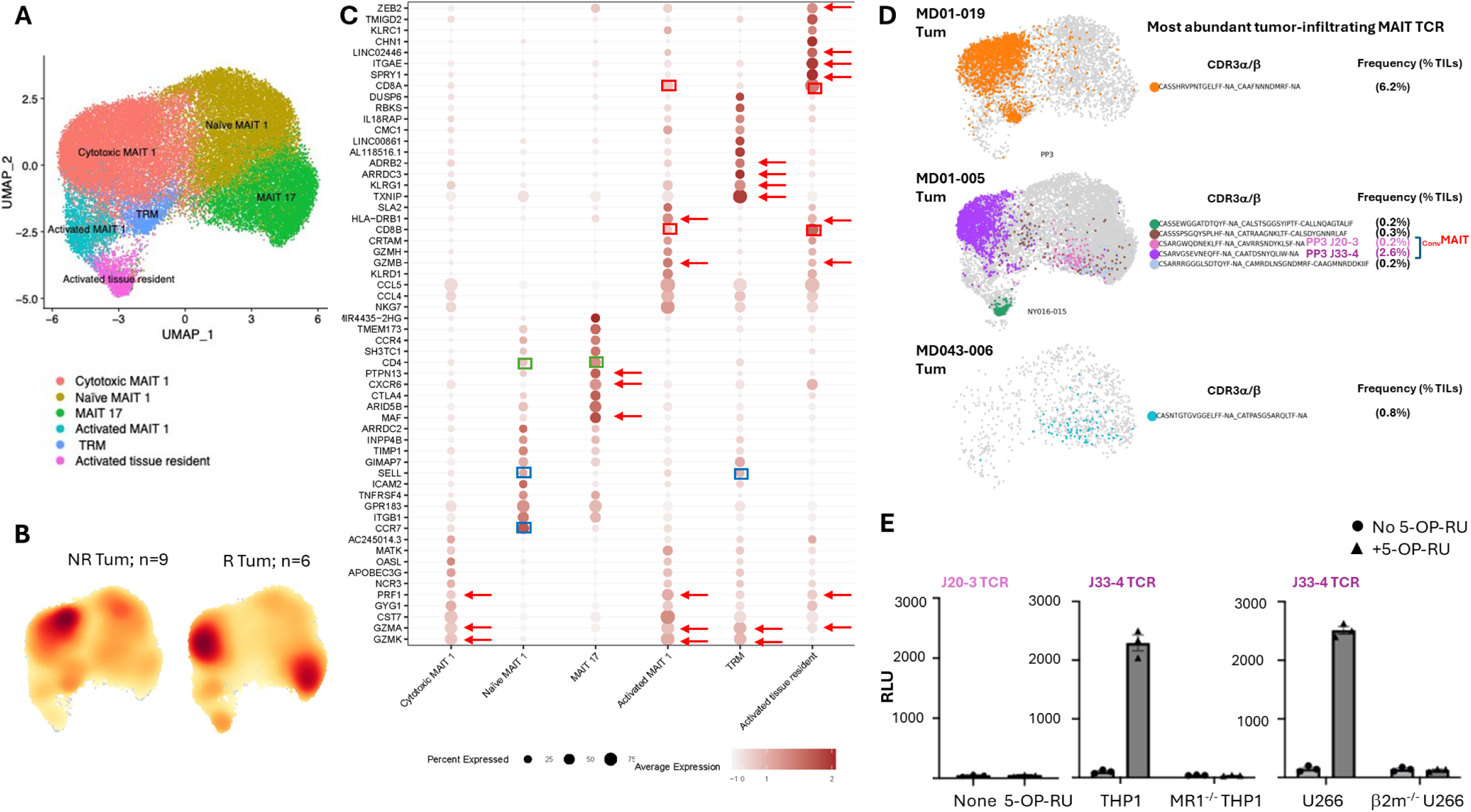
Single cell RNA and TCR sequencing analysis of intra-tumoral MAIT cells in lung cancer patients. **A,** UMAP representation of MAIT cell clusters resulting from the analysis of a published single cell RNA sequencing dataset (Caushi et al., 2021). **B**, Cell density plots of MAIT cells in tumor tissue separated by patient response status (Responders, R, n = 6; Non-responder, NR, NR= 9). **C**, Dot plot of top 10 markers for all 6 MAIT clusters. Red arrows indicate gene Id driving the cluster annotations. Red boxes indicate *CD8A* and *CD8B* genes, and green boxes indicate *CD4* gene. **D**, Overlay of the MAIT cell UMAP (shown in A) with single cell TCR sequencing indicates the 7 most prominent MAIT TCR clonotypes (color-coded), distributed across the 3 tumor samples, MD01-019, MD01-005 and MD043-006. The frequency of each clonotypes among the CD3+ TIL is indicated into brackets. _Conv_MAITs are noted according to the expression of the canonical recombination TCRAV1-2/J12,20,33. **E**, *Left,* MD01-005 J20-3 TCR from predicted _conv_MAIT cell did not recognize MR1 in the TCR capture assay in presence of 5-OP-RU; *Center*, MD01-005 J33-4 TCR from predicted _conv_MAIT cell recognized THP1 but not MR1-KO THP1 cells in presence of 5-OP-RU; *Right*, MD01-005 J33-4 TCR from predicted _conv_MAIT also recognized the human myeloma cell line, U266, but not b2M-KO U266 in presence of the 5-OP-RU.

We selected the predominant tumor-infiltrating _conv_MAIT clonotypes to carry bacteria recognition assays. The single cell RNA sequencing dataset was overlapped with the single cell TCR sequencing analysis (**Fig.1C**) to identify _conv_MAIT clonotypes (TCRVA 1-2(Germain et al., 2025)), which represented **7.5%** of MAITs. The two most frequent _conv_MAIT TCR clonotypes, *MAIT17* MD01-005 J20-3 (TCRAV1-2/J20; **0.2%** of the TIL) and *Cytotoxic MAIT1* MD01-005 J33-4 (TCRAV1-2/J33; **2.6%** of the TIL), were identified in R patient MD01-005 (**Fig.1C**). Although overall proportion of MAIT cells among T cells was similar between the R and NR patients, we found an increase of _conv_MAIT cells in R patients (16.7% v. 0.6% among MAIT cells in TILs of R and NR patients, respectively and representing 1.44% v. 0.06% among all T cells). However, this increase was primarily driven by the remarkable expansion of J33-4 MAIT clonotype in patient MD01-005.

MAIT TCR MD01-005 J20-3 and J33-4 TCRs were then cloned and validated for the MR1-dependent recognition of APC in presence of the riboflavin biosynthesis-associated metabolite, 5-OP-RU (**Fig.1D&E**). Our results showed that clonotypes MD01-005 J33-4 but not J20-3 MAIT TCR recognized 5-OP-RU-induced MR1, reinforcing the need to verify the MR1 dependence to functionally validate MAIT T cell properties. The most abundant _conv_MAIT TCR clonotype, MD01-005 J33-4, overlapping with *cytotoxic MAIT1* cluster was chosen to further probe how intra-tumoral bacteria impact the expression of MR1 on APC.

### Association between intratumoral lung cancer microbiota and _conv_MAIT cell infiltration

Since we postulated that local selective host-pathogen interactions orchestrate a tight regulation of MR1 translocation to cell surface membrane of target cells, avoiding inappropriate activation of MAIT cells, we first identified intratumoral microbiota by 16S rRNA amplicon sequencing of formalin-fixed paraffin embedded (FFPE) tumors (n=4) (**Fig.2A**).

**Figure 2.**
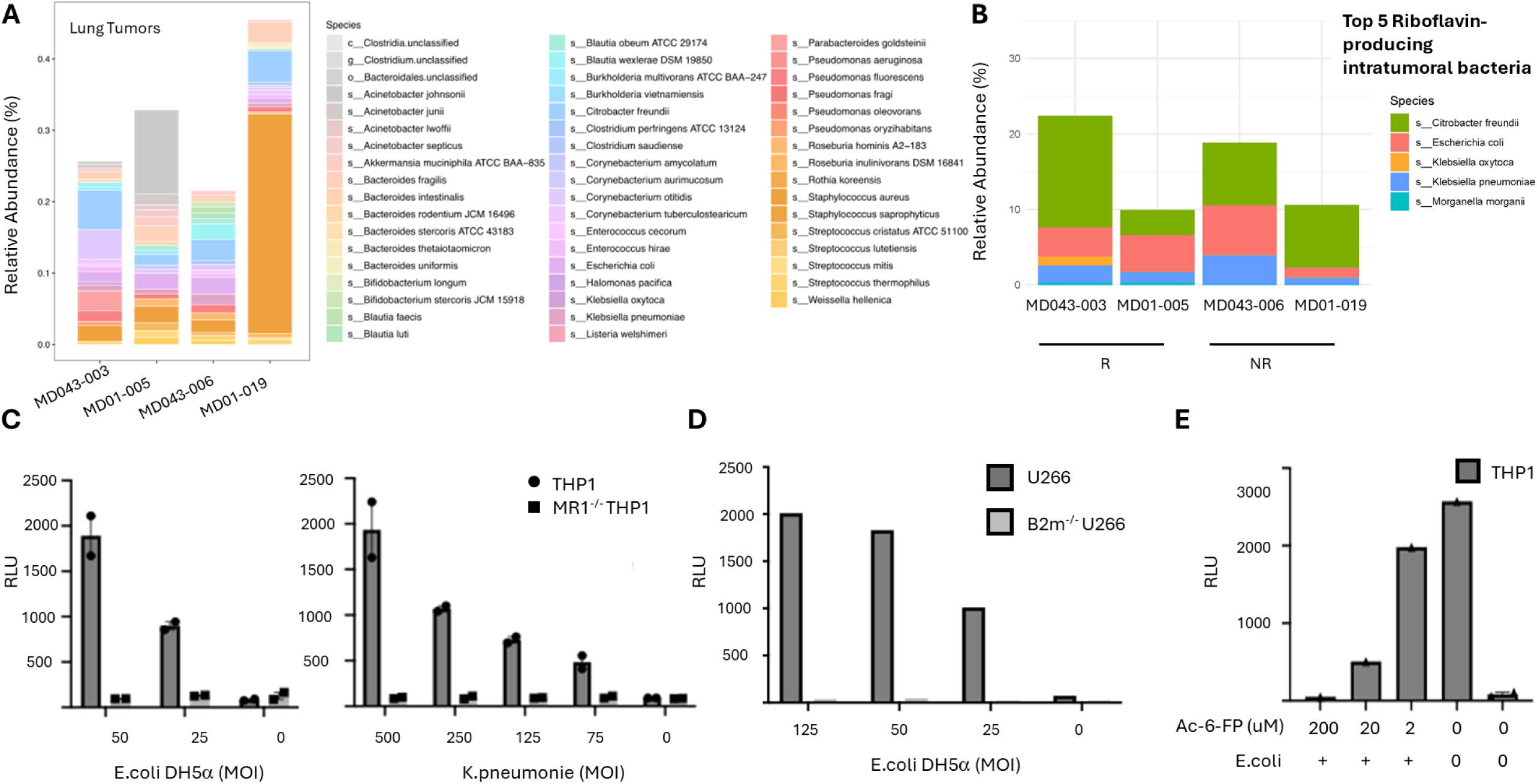
Intratumoral bacteria and Riboflavin metabolism detection in lung cancer. **A,** 16S V3-V4 RNA amplicon gene sequencing performed on DNA extracted from FFPE tumor tissue sections in four patients (MD01-019, MD01-005, MD043-006 and MD043-003). **B**, proportion (out of total bacteria) of the top 5 5-A-RU-producing intratumoral bacteria (Fig.S1) per patient. A literature-based pipeline was developed to predict the bacteria which will express at least the required three genes for 5-A-RU production (**Fig.S1B**). **C** Recognition of THP1 and MR1^-/-^ THP1 by J33-4 MAIT TCR expressing Jurkat reporter cells in presence of PFA-fixed *E. coli* DH5α or *K pneumonia* at different multiplicity of infection (MOI). Experiments were performed in replicates (n = 2) and are shown as mean ± SEM **D**, Recognition of U266 and b2m^-/-^ U266 by J33-4 MAIT TCR in presence of PFA-fixed *E. coli* DH5α. **E**, Inhibition of J33-4 MAIT TCR recognition of *E. coli* DH5α incubated THP1 by escalating Ac-6-FP doses (0.2, 20, 200 uM).

Next, to score how many of these bacteria had the ability to produce riboflavin-derived metabolites 5-A-RU, we developed a literature-based meta-analysis pipeline predicting MR1-binding metabolite biosynthesis ability of the tumor microbiome (**Fig.S1B-C**). We included a total of 3 genes core for 5-A-RU production: *ribA*, *ribE, yigB (PYRP2) (***Fig.S1B**)(Constantinides et al., 2019). We found that an average of 11% of the intratumoral bacteria in the 4 patients tested have the minimal three gene core required for 5-A-RU synthesis (**Fig.2B and Fig.S1C**). However, there was no association between predicted 5-A-RU biosynthesis and abundance of MAIT cells detected in our datasets (**Fig.S1D**). Noteworthy, we found that several strains of bacterial species previously reported to produce riboflavin like *Bacteroides thetaiotaomicron* and *Staphylococcus epidermidis* did not induce herein J33-4 MAIT TCR activation demonstrating that bacterial strains can vary in their ability to promote MAIT activation (**fig.S1E**).

We then confirmed that MD01-005’s J33-4 TCR recognized riboflavin-producing but not riboflavin-deficient bacteria in an MR1-dependent manner using PFA fixed *E. coli* and *K. pneumoniae* (both riboflavin synthesis proficient) versus riboflavin auxotrophic *E. hirae*. We showed that reporter jurkat cells expressing J33-4 TCR are activated by wild type (WT) THP1 or myeloma U266 cell lines, but not by their MR1 deficient counterparts (MR1^-/-^ THP1, B2m^-/-^ U266) in the presence of *Escherichia coli* or *Klebsiella pneumoniae (***Figure 2C&D**). Finally, we saw that J33-4 MAIT TCR activation is inhibited by escalating concentrations of the MR1 antagonistic ligand Acetyl 6 formyl pterin (Ac-6FP) (**Fig.2E**), which, although induces cell membrane expression of MR1 (**Fig.S1F**) is not recognized by MAIT TCR (**Fig.2E**) (Kjer-Nielsen et al., 2012). These data confirmed that patient J33-4 MAIT TCR activation is MR1 ligand dependent.

Altogether, our findings suggested that the identification of bacteria as well as the validation of the presence of the genes necessary to produce MR1 ligands are not sufficient to accurately predict MAIT TCR recognition. This may be, in part, because expression of the riboflavin precursor biosynthesis genes are tightly regulated by other environmental and contextual factors (Averianova et al., 2020). Furthermore, other metabolites such as folate can bind MR1 and block the activation of MAIT cells by riboflavin precursors (Kjer-Nielsen et al., 2012).

### Riboflavin-auxotrophic *Enterococcus spp,* but not *E. coli*, synergized with 5-OP-RU, to promote MR1 expression on APC, independently of pathogen associated membrane pattern (PAMP) recognition

Because bacteria can also produce antagonistic MR1 ligands (Kjer-Nielsen et al., 2012), we further tested whether *E. hirae* can suppress MAIT activation. While we did not have fresh tumor tissue available for microbiological analysis, we used clinical and ATCC available strains of species identified by 16S rRNA amplicon sequencing. Unexpectedly, rather than inhibiting MAIT TCR recognition, *E. hirae* triggered a synergistic effect on MAIT activation by 5-OP-RU, compared to 5-OP-RU alone (**Fig.3A**). Furthermore, we found that *E. faecalis*, *E. faecium* but not *E. avium,* exhibited the same synergistic effect. We confirmed by genomic sequencing and metabolomic analysis of the bacteria pellets that all *Enterococcus* spp used were riboflavin auxotrophic (**Fig.S2A&B**). We then showed that the synergistic properties of *Enterococcus* spp were not reproduced with other riboflavin auxotrophic bacteria tested (*B. longum*) and were selective to professional APC, including dendritic cells (DC), monocytic (THP1) and B cell-derived cell lines including human myeloma cell line U266 (**Fig.3B**). Human Melanoma (MeWo) and hepatocellular carcinoma (HCC) SNU387 cell lines, although capable of activating MAIT cells in presence of 5-OP-RU, did not exhibit higher MAIT activating properties in presence of *Enterococcus* (**Fig.3B**). We then verified by flow cytometry the cell surface expression of bioactive refolded MR1 using the 26.5 antibody (Huang et al., 2005). While surface expression of MR1 was previously reported as transient and low, **Fig.3C** shows that the membrane MR1 detection was not systematically matching TCR activation (Chua et al., 2011; Lamichhane and Ussher, 2017). In myeloma U266 cell line and human DC, but not THP1 for which MR1 was not detectable, higher activation of MAIT TCR by the combination 5-OP-RU/*E. faecalis* (**Fig.3A**) paralleled the increased MR1 ex-pression (**Fig.3C**). Suppression of synergic activation of J33-4/jurkat cells by increasing concentrations of ac-6FP provided a direct evidence of the role of increased MR1 cell surface expression in the synergy by *Enterococcus* (**Fig.3D**).

**Figure 3.**
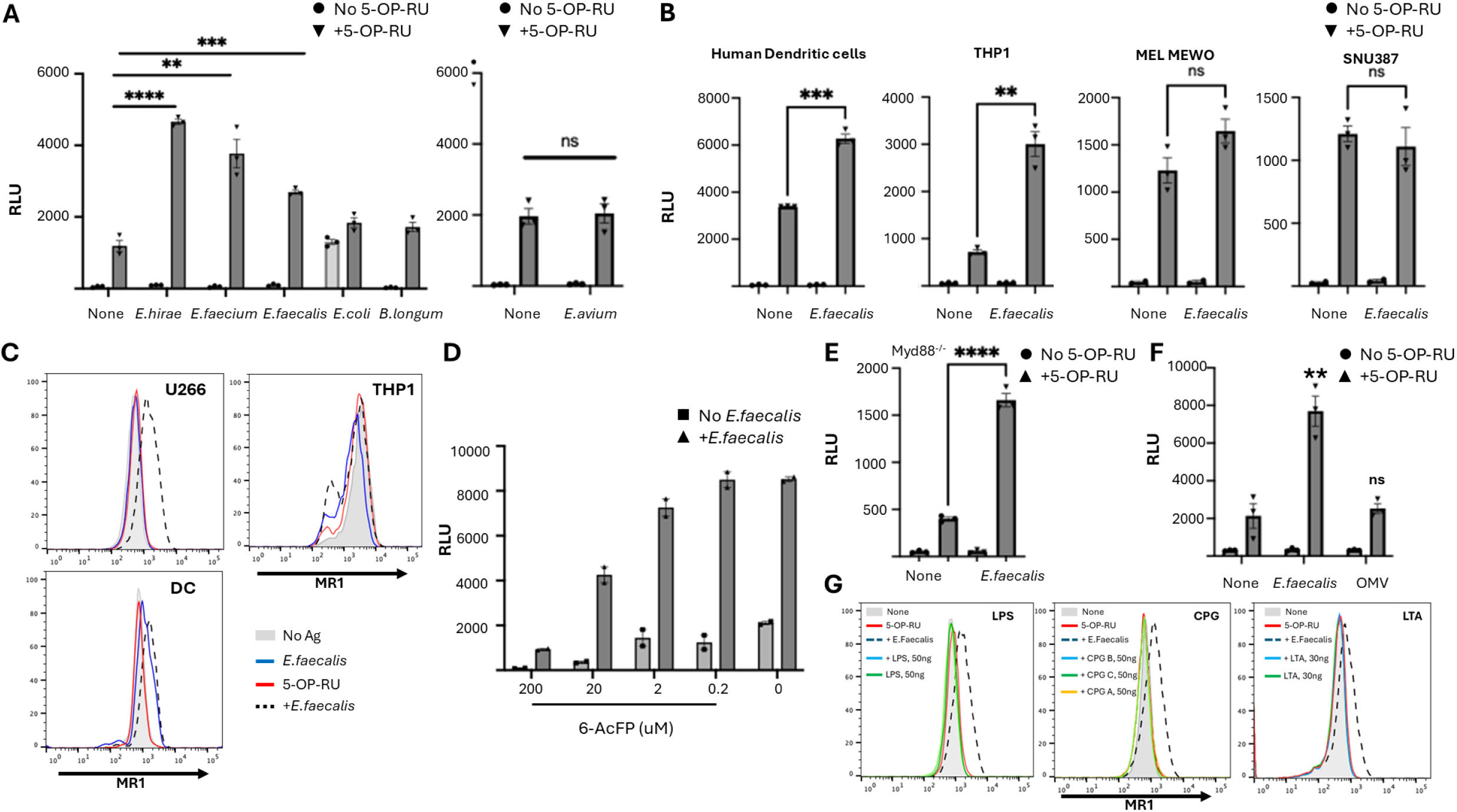
Synergistic role of Enterococcus in MAIT TCR activation by 5-OP-RU/MR1-presenting APC. **A,** J33-4 MAIT TCR recognition of U266 cells incubated overnight with (inverse black triangles) or without (black circles) 5-OP-RU, in presence or not of *E.hirae*, *E. faecalis*, *E. faecium*, *E. coli* and *B. longum* (Left graph) and *E. avium* (right graph, independent experiment). TCR activation is measured by bioluminescence reading. **B**, different APC (human dendritic cells, THP1, melanoma MeWO and hepatocellular carcinoma SNU387) were tested for the synergistic effect of *E faecalis* onto activation of conventional MAIT TCR J33-4 in presence (inverse black triangles) or not (black circles) of 5-OP-RU. **C**, Flow cytometry analysis of MR1 surface membrane expression using the 26.5 anti MR1 antibody staining of U266, THP1 and dendritic cells (DC) incubated overnight without or with 5-OP-RU in presence or not of *E.faecalis*. **D**, Inhibition by Ac-6-FP (escalating doses from 0.2 to 200 uM) of recognition by MAIT TCR J33-4 of MR1 expressed on U266 cultured or not with 5-OP-RU (6.25 uM) in presence (triangles) or not (squares) of *E.faecalis* (MOI:125). **E**, MAIT TCR J33-4 activation by Myd88^-/-^ THP1 cells incubated overnight with (inverse black triangles) or without (black circles) 5-OP-RU (6.25 uM), in presence or not of *E. faecalis (MOI:125).* **F**, MAIT TCR J33-4 activation by U266 cells incubated overnight with (inverse black triangles) or without (black circles) 5-OP-RU (6.25 uM), in presence or not of *E. faecalis (MOI:125) and E.faecalis* derived outer membrane vesicles (OMV). **G**, Flow cytometry analysis of MR1 surface membrane expression using the 26.5 anti MR1 antibody staining of U266 incubated overnight without or with 5-OP-RU in presence or not of *E. faecalis* (MOI:125), compared to LPS (TLR4 ligand; 50ng/ml; left histogram), CPG (TLR9 ligand; 5uM, middle histogram) or LTA (TLR2 ligand; 30ug/ml co; right histogram). All data are shown as mean ± SEM with technical replicates represented by number of symbols in each graph. ****:P<0.0001, ***:0.0001<P<0.001, **:0.001<P<0.01, *:0.01<P<0.05, ns (non significant) by unpaired t-test (two-tailed).

Culture of U266 with 5-OP-RU/*Enterococcus* did not increase the production of IL1β, IL-18 or IFNα in the supernatants compared to 5-OP-RU alone (**Fig.S2D**). Furthermore, the synergistic effect was observed only with live or PFA-fixed *Enterococcus spp*. Neither lysed nor sonicated bacteria triggered synergy (**Fig.S2E**). We therefore tested the possibility that *Enterococcus spp.* acted via PAMPs/pattern recognition receptor (PRR) interactions. However, Myd88-KO THP1 cells were as efficient as WT THP1 in activating J33-4 expressing Jurkat cells, (**Fig.3E**). Finally, neither non-replicative *E faecalis*-derived outer membrane vesicles (OVM, **Fig.3F**), or TLR ligands phenocopied the synergistic effect of *Enterococcus spp.* (**Fig.3G**). These results demonstrated that the synergistic effect of *Enterococcus spp*. was unlikely dependent on PAMP recognition. Interestingly, the synergistic effect of *Enterococcus spp.* was only observed with 5-OP-RU and not with PFA-fixed *E. coli*, indicating that the nature of the intracellular route of 5-OP-RU loaded MR1 (trans-golgi network [TGN] for soluble ligands versus phagolysosomes for bac-teria-derived MR1 ligand) was likely being impacted by *Enterococcus spp.* (**Fig S2F**).

### The synergistic effects of *Enterococcus spp.* on MR1-dependent MAIT cell activation is not transcriptionally regulated

The expression of MR1 and riboflavin precursors are ubiquitous but also key for the selective activation of MAIT cells (McWilliam et al., 2016). Previous studies showed that the expression of MR1 is highly contextual and diversely regulated according to the cell types [reviewed in (Karamooz et al., 2018) and (Germain et al., 2025)].

To understand the effect of *Enterococcus* infections onto MR1 up-regulation, we next performed bulk RNA sequencing of U266 cells incubated or not in presence of 5-OP-RU with *E. faecalis* (riboflavin auxotrophic) versus riboflavin-producing-E.coli (grp1=U266; grp2=U266/5-OP-RU; grp3=U266/Ef; grp4=U266/E.coli; grp5=U266/5-OP-RU+Ef; grp6=U266/5-OP-RU+E.coli **Fig.4**). We observed that although 5-OP-RU with *E. faecalis* led to higher expression of MR1 and activation of MAIT TCR compared to 5-OP-RU alone (**Fig.3C**), MR1 mRNA expression was not affected by *E. faecalis*, suggesting a post MR1 transcription synergistic effect of *E. faecalis*. Second, we observed that incubation of U266 with the combination 5-OP-RU/*E. faecalis* was associated with the upregulation of GO terms “*protein targeting to ER*” and KEGG signatures “*endocytosis*”, “*protein processing in ER”* compared to 5-OP-RU alone (**Fig. 4B&C**). Finally, compared with *E. coli*, *E. faecalis* co-culture led to the upregulation of a set of gene expression linked to autophagy and lipophagy such as DEPP1(Stepp et al., 2014), MT1G (Meng et al., 2024), BBS12(Pampliega et al., 2013), FICD(Gulen et al., 2024) and HILPDA(VandeKopple et al., 2019) (**Fig.S3**).

**Figure 4.**
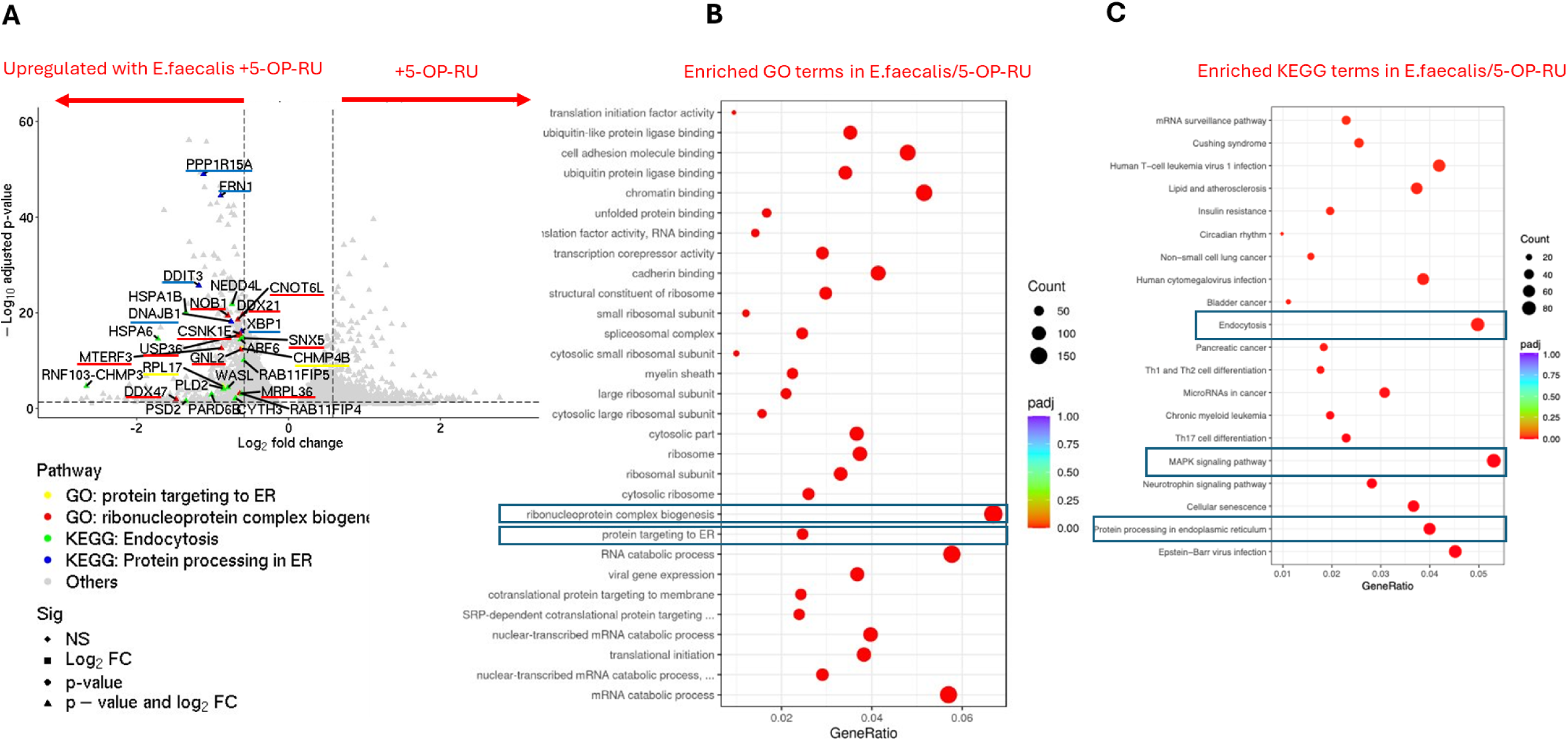
Differential effect of *Enterococcus faecalis* and *Escherichia coli* on gene expression by human myeloma U266 cells in presence or not of 5-OP-RU. **A**, volcano plot representing RNAseq analysis of U266 cell line incubated with 5-OP-RU + *E.faecalis* compared to 5-OP-RU alone [p<0.01 and −Log2(FC>2)]. Genes belonging to the GO signature “*protein targeted to ER*” and “*ribonucleoprotein complex biogenesis*” are annotated in yellow and red respectively. Genes included in the KEGG signature “*protein targeting to ER*” and “*Endocytosis*” are annotated in blue and red respectively. **B&C**, Dot plot visualization of the enriched GO (**B**) and KEGG (**C**) terms in 5-OP-RU/E. faecalis condition after RNAseq analysis of U266 cells culture in 5-OP-RU + *E faecalis* compared to 5-OP-RU alone.

Altogether, these results suggest that Enterococcus may rather perturb vacuolar trafficking and MR1 translocation to the cytoplasmic membrane.

### Synergistic activity of *Enterococcus* is mediated by alteration of MR1 intracellular traffick-ing

*E. faecalis* is capable of persisting in the endo-lysosomal compartment and resisting autophagy in macrophages via inhibition of the fusion of lysosome with late endosomes or autophagosomes, respectively (da Silva et al., 2022; Zou and Shankar, 2016). We therefore tested the role of autophagy and intracellular endo-lysosomal trafficking pathway in the regulation of MR1 cell expression at the cell surface of APC by *Enterococcus*. We used a set of pharmacological agents which induce (Rapamycin; RAPA) or inhibit (*Wortmannin*, or WTM, an inhibitor of the PI3K, Cytochalasin D or CYTO D, an inhibitor of actin polymerization) autophagy, inhibits TGN trafficking (Brefeldin A; BFA), *de novo* protein synthesis (Cycloheximide; CHX) or the endo-lysosome pathway (Bafilomycin A, or Baf A1, an inhibitor of vacuolar type-H+ ATPase and Chloroquine or CHQ, both inhibiting the endosome acidification) (**Fig.5A**). Importantly, upon dose optimization, none of these pharmacological inhibitors were cytotoxic for the APC (>80% viability).

**Figure 5.**
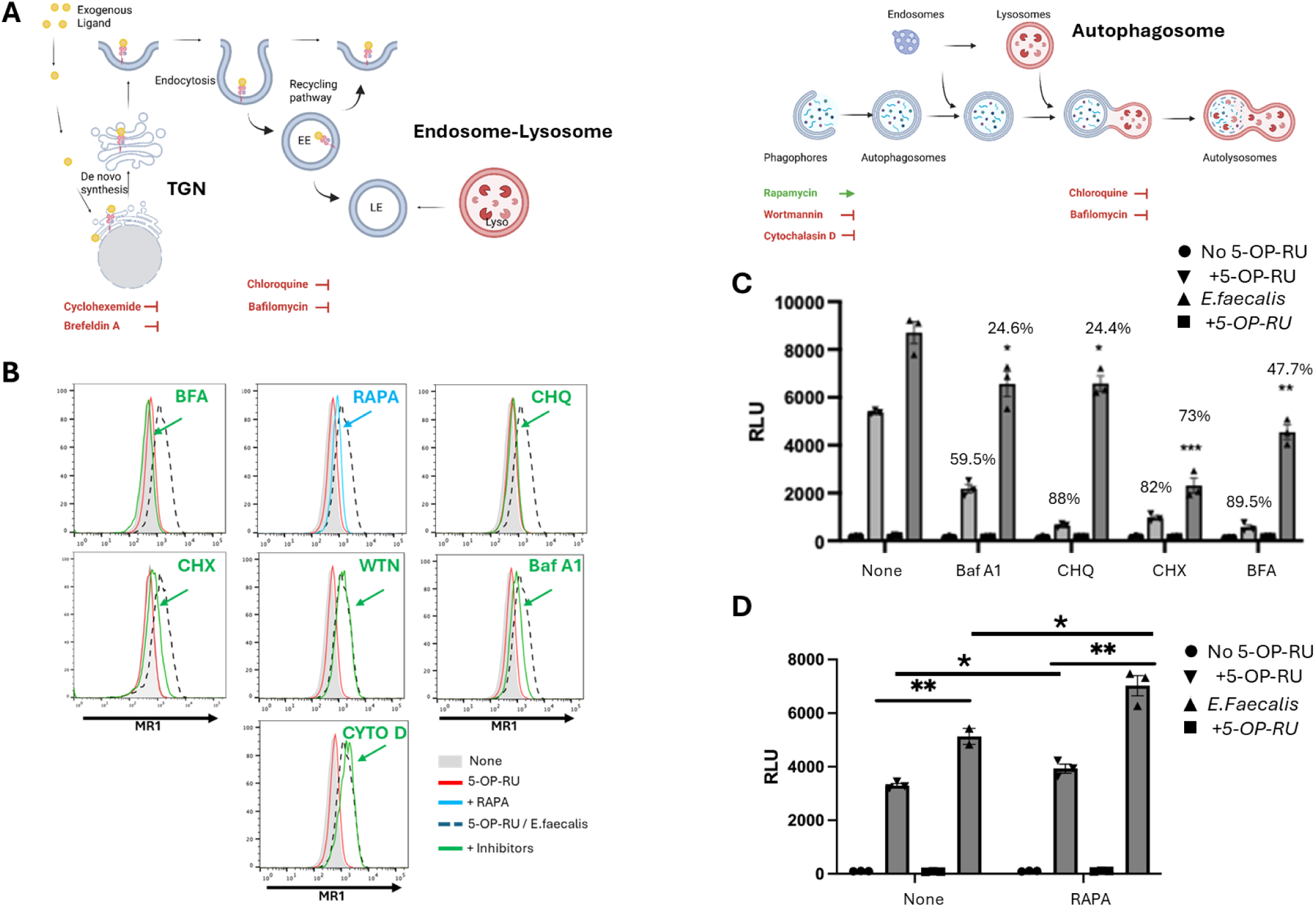
*Enterococcus spp* interfered with MR1 endo-lysosomal trafficking and cell surface translocation. **A**, schematic representation of MR1 intra-cellular translocation along the transgolgi network (TGN) to the cell surface membrane, the endosome-lysosome and the autophagolysome formation pathways. The different pharmacological inhibitors (in red; BFA, CHX, Baf A1, CHQ, CYTO D, WTN) of intracellular trafficking and an inducer of autophagy (RAPA in blue) used during the incubation of U266 with 5-OP-RU (6.25uM) in presence or not of *E.faecalis* (MOI:125) are indicated below the pathway they are targeting. **B**, Flow cytometry analysis of MR1 expression by U266 using the 26.5 antibody clone after overnight incubation with 5-OP-RU and *E. faecalis* in presence of Baf A1 (100nM), ChQ (100uM), CHX (20ug/ml), BFA (3ug/ml) and RAPA (100nM). **C & D**, 6 hr-MR1 recognition assay using J33-4 MAIT TCR expressing jurkat reporter cells and, as APC, U266 treated overnight with the drugs described in **B** (APC were washed after incubations with pharmacological agents). Results are shown as relative values of TCR/NFAT-induced bioluminescence (RLU). Schemas realized with Biorender(R) graphics. All data are shown as mean ± SEM with technical replicates represented by number of symbols in each graph. ****:P<0.0001, ***:0.0001<P<0.001, **:0.001<P<0.01, *:0.01<P<0.05, ns (non significant) by unpaired t-test (two-tailed).

First, while CHQ and Baf A1 (endo-lysosome pathway) partially reduced MR1 expression on U266 cultured in presence of *E. faecalis* and 5-OP-RU, BFA and CHX (secretory pathway) had a more pronounced effect on reducing cell surface MR1 (**Fig.5B**). This suggested that, while the trafficking of MR1 through TGN (BFA and CHX sensitive) predominantly fuels the pool of surface membrane-bound MR1, a fraction of endocytosed MR1 can be recycled from the early endosomal compartment (CHQ and Baf A1 sensitive) back to the cell surface, instead of being degraded in late endosomes (**Fig.5B**). In parallel, **Fig. 5C** shows that the activation of MAIT TCR J33-4 was ablated after overnight incubation of APC with BFA and CHX (**89.5%** and **82%** inhibition respectively), confirming that MR1 cell surface expression (**Fig.5C**) and TCR recognition required *de novo* MR1 synthesis and transport from the ER to the Golgi complex. Previous studies have already established that exogenous ligand can be loaded on MR1 in the ER and trigger the translocation of MR1 to the surface (McWilliam et al., 2016). Our data further showed that the activation of MAIT cells was partially inhibited by Baf A1 (**59.4%**) but much more strongly by less specific CHQ (**88%**), again confirming our flow cytometry data and supporting previous reports that at least a fraction of MR1 resides in late endosomes where they cargo endogenous ligands before reaching the cell surface and activate MAIT cells (recycling MR1). Subsequently adding *E. faecalis* to overnight incubation of U266 with 5-OP-RU, radically modified the sensitivity of MR1 cell surface translocation and MAIT TCR recognition to the set of inhibitors tested (**Fg.5B-D**) confirming that, overall, *Enterococcus spp* perturb MR1 intr-cellular trafficking. Whereas MAIT TCR recognition was again predominantly dependent on *de novo* synthesis of MR1 (**73%** inhibition MAIT activation by CHX), they became only partially dependent on TGN trafficking (**47%** inhibition by BFA) and marginally inhibited by CHQ and Baf A1 (**24.4%** and **24.6%** inhibition, respectively) in presence of *E. faecalis* (**Fig.5C**). This may indicate that alteration of the late endosome / lysosome pathway by *E. faecalis* (da Silva et al., 2022) may boost the accumulation of MR1 in early endosome compartment via limitation of MR1 lysosomal degradation rate and in return, overflow of MR1 recycling to the plasma membrane (**Fig.6**). The increased density of MR1 at the cell surface of APC is expected to be locally sensed by and activate MAIT cells. Hariff’s group previously showed that the regulation of MR1 expression is primarily tuned by pre-synthesized MR1 intra-cellular trafficking from MR1 vesicles (ER-like subcellular compartment) but also in certain circumstances early endosomal-dependent recycling pathway (Harriff et al., 2016; Huber et al., 2020). Finally, although CYTO D and WTN did not alter MR1 cell surface expression upon 5-OP-RU/*E. faecalis* incubation, RAPA partially increased the expression of MR1 cultured with 5-OP-RU (**Fig.5B**) and MAIT TCR recognition (**Fig.5D**), suggesting that autophagy may contribute, at least partially, to the synergistic upregulation of MR1.

**Figure 6.**
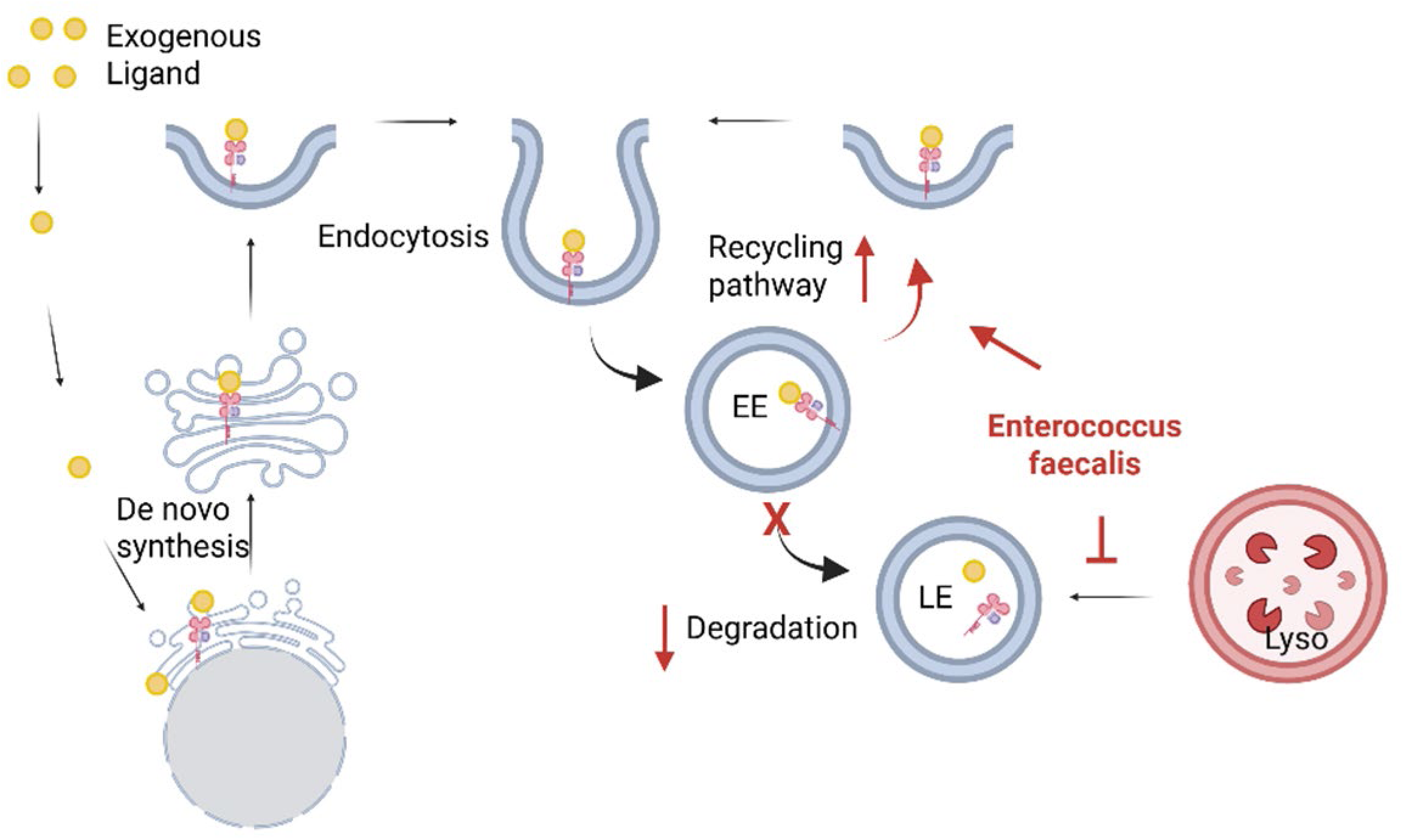
Schematic representation of the mechanism used by innate immunity to detect intracellular *Enterococcus spp*. *Enterococcus spp* blocked endo-lysosomal degradation pathways forcing an increased early endosomal MR1 recycling to the cell surface membrane. Shema realized with Biorender(R) graphics.

### Concluding remark

Gut microbiota has emerged in these past decades as a potent actor in the efficacy of cancer treatment, especially immune checkpoint inhibitors (Elkrief et al., 2025; Griffin and Hang, 2022). However, little is known about the molecular mechanisms by which immune modulatory microbes may act on cancer immunosurveillance and response to immunotherapy. We herein, unveil a possible innate mechanism by which intratumoral bacteria altering the invariant immune ligands density and cell surface turn over may be sensed as a new form of intracellular bacteria-associated pattern, promoting the stimulation and expansion of innate-like T cells. Although riboflavin-auxotrophic *Enterococcus spp*, detected in the tumor of lung cancer patients, are not capable of directly stimulating MAIT cells, they demonstrated their synergistic properties in enhancing the expression of ligand-loaded MR1 at the surface of APC, including DCs, monocytic phagocytes and B cells. Therefore, certain intracellular pathogens may be sensed by the immune system through modification of the MR1 recycling rate and its accumulation at the surface of infected cells. Several *Enterococcus* species including *hirae*, *faecalis* or *faecium* were enriched in the gut of ICB responsive cancer patients and preclinical models showed their role in enhancing cancer immunotherapy (Griffin et al., 2021). Although we couldn’t conclude regarding the *in vivo* role of MAIT cells in the pathological response to ICI, this result may open the perspective of adjuvant probiotic bacterial components enhancing the response to ICI *via* boosting intratumoral innate immunity.

### Weaknesses of the study

The scope of the study is limited to the role of *Enterococcus* in regulating of MR1 expression for TCR recognition. The limited number of responding patients precluded correlative studies which could suggest the role of MAIT cells in pathological responses to PD-1 blockade. Finally, the study does not clarify whether *Enterococcus* species enter the target cells or remain extracellular to mediate their effects.

## Materials and Methods

### Lung cancer patient single cell dataset and MAIT cell cluster analysis

Single cell RNA and TCR sequencing data from patients with NSCLC treated with neoadjuvant ICB was obtained from previously published dataset (Caushi et al., 2021). Clustering of the annotated CD3+ MAIT cell cluster was performed using Seurat V5. Dimensional reduction was done using the RunUMAP function. Cell markers were identified by using a two-sided Wilcoxon rank sum test. Genes with adjusted *P< 0.05* were retained. MAIT sub-clusters were annotated based on differential genes and immune cell markers. Conventional MAIT cells defined by the presence of TRAV1-2 gene were overlaid on MAIT cell population.

### Cell lines and reagents

Genetically modified CD8-expressing Jurkat reporter cell line, CD8+NFAT-lucΔ𝛼𝛼𝛼𝛼, was kindly provided by Dr Smith(Caushi et al., 2021). U266 cells were kindly provided by Dr I. Borello’s lab and cultured in RPMI with 10% FBS, 200uM glutamine and 1% pen-strep. MeWo and SNU387 cell lines were kindly provided by Dr. Scott Wilson’s lab and cultured according to ATCC protocol. THP1 cells were obtained from ATCC and cultured following the provider’s recommendation. Beta-2 microglobulin-knockout (𝛼𝛼2m KO) Jurkat cells and 𝛼𝛼2m KO U266 cells were generated via CRISPR-Cas9 system synthetic guide RNA (sgRNA) targeting: GAGTAGCGCGAGCACAGCTAAGG (Synthego, Redwood City, CA). An MR1-KOTHP1 cell line was generated via CRISPR-Cas9 with sgRNA targeting: TGGAACTGAAGCGCCTACAGAGG.

5-A-RU-PABC-Val-Cit-Fmoc is a prodrug of the MR1 ligand 5-A-RU, a precursor of Riboflavin (Vitamin B2) and purchased at MedchemExpress (Monmouth Junction, NJ). Brefeldin A and LPS were purchased from Thermofisher (Waltham, MA). Chloroquine was purchased from Biolegend (SanDiego, CA). Cycloheximide, Cytochalasin D, Bafilomycin A, Rapamycin, Wortmannin, and Lipoteichoic acid (LTA) were purchased from Millipore Sigma (Saint Louis, MI). TLR9 was purchased at Invivogen (SanDiego, CA).

### Bacteria

A set of bacteria Robiflavin producing or deficient) was cultured in medium described in Table 1 and aliquoted at defined colony-forming units (CFU). Clinical isolates were kindly provided by Dr C Sears.

### Bacteria strains used in the TCR capture functional assay

| **Bacteria** | **Strain** | **Medium**<br>**(broth / plate)** | **Oxygen** |
| --- | --- | --- | --- |
| *Escherichia coli* | DH5α | LB / LB | Aerobic |
| *Klebsiella pneumoniae* | Clinical isolate | BHI / BHI | Aerobic |
| *Enterococcus spp* | Clinical isolate | BHI / BHI | Aerobic |
| *Pseudomonas aeruginosa* | Clinical isolate | TSB / TSA | Aerobic |
| *Staphylococcus aureus* | SH1000 | TSB/ TSA blood | Aerobic |
| *Staphylococcus epidermidis* | Clinical isolate | TSB / TSA | Aerobic |
| *Bacteriodes thetaiotaomicron* | VP5482 | BHI /BHI | Anaerobic |
| *Bifidobacterrium longum* | ATCC15708 | RC / BHI | Anaerobic |

To prepare bacteria for use in MAIT cell assays, cultures were centrifuged at 4,000g for 15min. Bacterial pellets were resuspended in 50% glycerol:culture media and stored at −80℃ in single-use aliquot. Prior to use with mammalian cells in most assays, bacteria were fixed using 2% PFA for 15min. Additional methods tested include freeze-thaw cycles (20 cycles liquid nitrogen / room temperature), heat inactivation (70℃ for 30 minutes), and sonication (*Q-SONICA* sonicator at 50 amplitude for 20 cycles of 10 sec-sonication and 1 min-rest on ice). For live bacteria pre-incubation, experimental aliquots were washed vigorously with PBS and incubated with cell lines at 37℃ for different times, followed by vigorous wash with PBS (277g, 5min per time).

### TCR transfection and transient expression

The protocol was adapted from previously described methods(Caushi et al., 2021). Briefly, we identified TRAV and TRBV families using the IMGT/V-Quest database (http://www.imgt.org) and the TCR were generated by fusing the TCRA V-J region sequences and the TCRB V-D-J regions with the human TRA and TRB constant chains, respectively. The full-length TCRA and TCRB chains were then produced as individual gene blocks and cloned into a pCI mammalian expression vector, containing a CMV promoter followed by transformation of competent NEB® 5-alpha Competent *E. coli (New England Biolabs*) following manufacturer protocol*. The integrated plasmid DNA was extracted via QIAprep Spin Miniprep Kit* and stored at −20℃. DNA plasmid was checked by Sanger sequencing (Genewiz). Jurkat reporter cells were then electroporated with pCI plasmid encoding TCRa and TCRb genes using ECM830 Square wave electroporation system (BTX) at 275 V for 10 ms in OptiMem media in a 4-mm cuvette. Post electroporation, cells were rested overnight by incubating in RPMI 10% FBS at 37 °C, 5% CO_2_. TCR expression was confirmed by flow cytometric staining for CD3 ex-pression.

### In vitro MR1 recognition assay and flow cytometry

1.5 × 10^5^ CD3+ TCR-expressing CD8+NFAT-lucΔ𝛼𝛼𝛼𝛼 reporter jurkat cells were co-cultured with 1.5 × 10^5^ target cells (WT THP1 and MR1 KO THP1) in the presence of pro-drug 5-A-RU. Anti-CD3 (OKT3 clone) was used as a positive control. After overnight incubation, activation of the NFAT reporter gene was measured by the Bio-Glo Luciferase Assay following the manufacturer’s instructions (Promega). In some experiments APC were preincubated overnight with BFA (3ug/ml), CHX (20ug/ml), CHQ (100uM), Bafilomycin A1 (100nM) and Rapamycin (100nM) in presence of 6.25nM of 5-OP-RU with *Enterococcus* spp or vehicle control. Cells were then washed, and one aliquot was used to test MR1 surface expression by flow cytometry with anti-MR1 antibody 26.5 clone (Biolegend). The other aliquot was used to test TCR recognition as described above.

### Bulk RNA sequencing

U266 myeloma cell line was cultured or not (untreated control) in presence of Riboflavin in presence or not of *E. coli* versus *E. faecalis* overnight. RNA was extracting using MiniRNA kit (Qiagen; Germantown, MD) and bulk RNAseq was performed by Novogene (Sacremento, CA).

### Metabolomic

Pellets and supernatants of bacteria culture were analysed for their content in riboflavin and folate using ML-MS / MS approach by *Creative Proteomics* (Shirley, NY)

### ELISA analysis

IFNα, IL1β and TNFα were measured in the overnight U266 culture supernatants using R&D System ELISA kits following the manufacturer’s protocols.

### Statistics

All statistical analyses were performed utilizing Graphpad Prism v10 (excluding single RNA sequencing and bulk RNA sequencing analysis). A p-value of less than 0.05 was considered statistically significant. Unpaired two-tailed t-test was used for statistical comparisons as indicated in the figure legends. P value legend is as follows- ****:P<0.0001, ***:0.0001<P<0.001, **:0.001<P<0.01, *:0.01<P<0.05, ns (non significant) by unpaired t-test (two-tailed).

## REFERENCES

1. Averianova, L.A., L.A. Balabanova, O.M. Son, A.B. Podvolotskaya, and L.A. Tekutyeva. 2020. Production of Vitamin B2 (Riboflavin) by Microorganisms: An Overview. Front Bioeng Biotechnol 8:570828.

2. Bugaut, H., Y. El Morr, M. Mestdagh, A. Darbois, R.A. Paiva, M. Salou, L. Perrin, M. Furstenheim, A. du Halgouet, L. Bilonda-Mutala, A.L. Le Gac, M. Arnaud, A. El Marjou, C. Guerin, A. Chaiyasitdhi, J. Piquet, D.M. Smadja, A. Cieslak, B. Ryffel, V. Maciulyte, J.M.A. Turner, K. Bernardeau, X. Montagutelli, O. Lantz, and F. Legoux. 2024. A conserved transcriptional program for MAIT cells across mammalian evolution. J Exp Med 221:

3. Caushi, J.X., J. Zhang, Z. Ji, A. Vaghasia, B. Zhang, E.H. Hsiue, B.J. Mog, W. Hou, S. Justesen, R. Blosser, A. Tam, V. Anagnostou, T.R. Cottrell, H. Guo, H.Y. Chan, D. Singh, S. Thapa, A.G. Dykema, P. Burman, B. Choudhury, L. Aparicio, L.S. Cheung, M. Lanis, Z. Belcaid, M. El Asmar, P.B. Illei, R. Wang, J. Meyers, K. Schuebel, A. Gupta, A. Skaist, S. Wheelan, J. Naidoo, K.A. Marrone, M. Brock, J. Ha, E.L. Bush, B.J. Park, M. Bott, D.R. Jones, J.E. Reuss, V.E. Velculescu, J.E. Chaft, K.W. Kinzler, S. Zhou, B. Vogelstein, J.M. Taube, M.D. Hellmann, J.R. Brahmer, T. Merghoub, P.M. Forde, S. Yegnasubramanian, H. Ji, D.M. Pardoll, and K.N. Smith. 2021. Transcriptional programs of neoantigen-specific TIL in anti-PD-1-treated lung cancers. Nature 596:126–132.

4. Chua, W.J., S. Kim, N. Myers, S. Huang, L. Yu, D.H. Fremont, M.S. Diamond, and T.H. Hansen. 2011. Endogenous MHC-related protein 1 is transiently expressed on the plasma membrane in a conformation that activates mucosal-associated invariant T cells. J Immunol 186:4744–4750.

5. Constantinides, M.G., V.M. Link, S. Tamoutounour, A.C. Wong, P.J. Perez-Chaparro, S.J. Han, Y.E. Chen, K. Li, S. Farhat, A. Weckel, S.R. Krishnamurthy, I. Vujkovic-Cvijin, J.L. Linehan, N. Bouladoux, E.D. Merrill, S. Roy, D.J. Cua, E.J. Adams, A. Bhandoola, T.C. Scharschmidt, J. Aube, M.A. Fischbach, and Y. Belkaid. 2019. MAIT cells are imprinted by the microbiota in early life and promote tissue repair. Science 366:

6. Corbett, A.J., S.B. Eckle, R.W. Birkinshaw, L. Liu, O. Patel, J. Mahony, Z. Chen, R. Reantragoon, B. Meehan, H. Cao, N.A. Williamson, R.A. Strugnell, D. Van Sinderen, J.Y. Mak, D.P. Fairlie, L. Kjer-Nielsen, J. Rossjohn, and J. McCluskey. 2014. T-cell activation by transitory neo-antigens derived from distinct microbial pathways. Nature 509:361–365.

7. Crowther, M.D., G. Dolton, M. Legut, M.E. Caillaud, A. Lloyd, M. Attaf, S.A.E. Galloway, C. Rius, C.P. Farrell, B. Szomolay, A. Ager, A.L. Parker, A. Fuller, M. Donia, J. McCluskey, J. Rossjohn, I.M. Svane, J.D. Phillips, and A.K. Sewell. 2020. Genome-wide CRISPR-Cas9 screening reveals ubiquitous T cell cancer targeting via the monomorphic MHC class I-related protein MR1. Nat Immunol 21:178–185.

8. da Silva, R.A.G., W.H. Tay, F.K. Ho, F.R. Tanoto, K.K.L. Chong, P.Y. Choo, A. Ludwig, and K.A. Kline. 2022. Enterococcus faecalis alters endo-lysosomal trafficking to replicate and persist within mammalian cells. PLoS Pathog 18:e1010434.

9. Dolton, G., H. Thomas, L.R. Tan, C. Rius Rafael, S. Doetsch, G.A. Ionescu, L.F. Cardo, M.D. Crowther, E. Behiry, T. Morin, M.E. Caillaud, D. Srai, L. Parolini, M.S. Hasan, A. Fuller, K. Topley, A. Wall, J.R. Hopkins, N. Omidvar, C. Alvares, J. Zabkiewicz, J. Frater, B. Szomolay, and A.K. Sewell. 2025. MHC-related protein 1-restricted recognition of cancer via a semi-invariant TCR-alpha chain. J Clin Invest 135:

10. Dominguez, R., L.W. Young, J. Ledesma-Medina, J. Cienfuegos, J.C. Gartner, K.M. Bron, and T.E. Starzl. 1985. Pediatric liver transplantation. Part II. Diagnostic imaging in postoperative management. Radiology 157:339–344.

11. Elkrief, A., R. Pidgeon, S. Maleki Vareki, M. Messaoudene, B. Castagner, and B. Routy. 2025. The gut microbiome as a target in cancer immunotherapy: opportunities and challenges for drug development. Nat Rev Drug Discov 24:685–704.

12. Germain, L., P. Veloso, O. Lantz, and F. Legoux. 2025. MAIT cells: Conserved watchers on the wall. J Exp Med 222:

13. Gherardin, N.A., A.N. Keller, R.E. Woolley, J. Le Nours, D.S. Ritchie, P.J. Neeson, R.W. Birkinshaw, S.B.G. Eckle, J.N. Waddington, L. Liu, D.P. Fairlie, A.P. Uldrich, D.G. Pellicci, J. McCluskey, D.I. Godfrey, and J. Rossjohn. 2016. Diversity of T Cells Restricted by the MHC Class I-Related Molecule MR1 Facilitates Differential Antigen Recognition. Immunity 44:32–45.

14. Griffin, M.E., J. Espinosa, J.L. Becker, J.D. Luo, T.S. Carroll, J.K. Jha, G.R. Fanger, and H.C. Hang. 2021. Enterococcus peptidoglycan remodeling promotes checkpoint inhibitor cancer immunotherapy. Science 373:1040–1046.

15. Griffin, M.E., and H.C. Hang. 2022. Microbial mechanisms to improve immune checkpoint blockade responsiveness. Neoplasia 31:100818.

16. Gulen, B., A. Blevins, L.N. Kinch, K.A. Servage, N.M. Stewart, H.F. Gray, A.K. Casey, and K. Orth. 2024. FicD sensitizes cellular response to glucose fluctuations in mouse embryonic fibroblasts. Proc Natl Acad Sci U S A 121:e2400781121.

17. Harriff, M.J., E. Karamooz, A. Burr, W.F. Grant, E.T. Canfield, M.L. Sorensen, L.F. Moita, and D.M. Lewinsohn. 2016. Endosomal MR1 Trafficking Plays a Key Role in Presentation of Mycobacterium tuberculosis Ligands to MAIT Cells. PLoS Pathog 12:e1005524.

18. Huang, S., S. Gilfillan, M. Cella, M.J. Miley, O. Lantz, L. Lybarger, D.H. Fremont, and T.H. Hansen. 2005. Evidence for MR1 antigen presentation to mucosal-associated invariant T cells. J Biol Chem 280:21183–21193.

19. Huber, M.E., R. Kurapova, C.M. Heisler, E. Karamooz, F.G. Tafesse, and M.J. Harriff. 2020. Rab6 regulates recycling and retrograde trafficking of MR1 molecules. Sci Rep 10:20778.

20. Ito, E., S. Inuki, Y. Izumi, M. Takahashi, Y. Dambayashi, L. Ciacchi, W. Awad, A. Takeyama, K. Shibata, S. Mori, J.Y.W. Mak, D.P. Fairlie, T. Bamba, E. Ishikawa, M. Nagae, J. Rossjohn, and S. Yamasaki. 2024. Sulfated bile acid is a host-derived ligand for MAIT cells. Sci Immunol 9:eade6924.

21. Karamooz, E., M.J. Harriff, and D.M. Lewinsohn. 2018. MR1-dependent antigen presentation. Semin Cell Dev Biol 84:58–64.

22. Keller, A.N., S.B. Eckle, W. Xu, L. Liu, V.A. Hughes, J.Y. Mak, B.S. Meehan, T. Pediongco, R.W. Birkinshaw, Z. Chen, H. Wang, C. D’Souza, L. Kjer-Nielsen, N.A. Gherardin, D.I. Godfrey, L. Kostenko, A.J. Corbett, A.W. Purcell, D.P. Fairlie, J. McCluskey, and J. Rossjohn. 2017. Drugs and drug-like molecules can modulate the function of mucosal-associated invariant T cells. Nat Immunol 18:402–411.

23. Kjer-Nielsen, L., O. Patel, A.J. Corbett, J. Le Nours, B. Meehan, L. Liu, M. Bhati, Z. Chen, L. Kostenko, R. Reantragoon, N.A. Williamson, A.W. Purcell, N.L. Dudek, M.J. McConville, R.A. O’Hair, G.N. Khairallah, D.I. Godfrey, D.P. Fairlie, J. Rossjohn, and J. McCluskey. 2012. MR1 presents microbial vitamin B metabolites to MAIT cells. Nature 491:717–723.

24. Koay, H.F., N.A. Gherardin, C. Xu, R. Seneviratna, Z. Zhao, Z. Chen, D.P. Fairlie, J. McCluskey, D.G. Pellicci, A.P. Uldrich, and D.I. Godfrey. 2019. Diverse MR1-restricted T cells in mice and humans. Nat Commun 10:2243.

25. Lamichhane, R., and J.E. Ussher. 2017. Expression and trafficking of MR1. Immunology 151:270–279.

26. Le Bourhis, L., M. Dusseaux, A. Bohineust, S. Bessoles, E. Martin, V. Premel, M. Core, D. Sleurs, N.E. Serriari, E. Treiner, C. Hivroz, P. Sansonetti, M.L. Gougeon, C. Soudais, and O. Lantz. 2013. MAIT cells detect and efficiently lyse bacterially-infected epithelial cells. PLoS Pathog 9:e1003681.

27. Le Bourhis, L., E. Martin, I. Peguillet, A. Guihot, N. Froux, M. Core, E. Levy, M. Dusseaux, V. Meyssonnier, V. Premel, C. Ngo, B. Riteau, L. Duban, D. Robert, S. Huang, M. Rottman, C. Soudais, and O. Lantz. 2010. Antimicrobial activity of mucosal-associated invariant T cells. Nat Immunol 11:701–708.

28. Lepore, M., A. Kalinichenko, S. Calogero, P. Kumar, B. Paleja, M. Schmaler, V. Narang, F. Zolezzi, M. Poidinger, L. Mori, and G. De Libero. 2017. Functionally diverse human T cells recognize non-microbial antigens presented by MR1. Elife 6:

29. Malenica, I., J. Adam, S. Corgnac, L. Mezquita, E. Auclin, I. Damei, L. Grynszpan, G. Gros, V. de Montpreville, D. Planchard, N. Theret, B. Besse, and F. Mami-Chouaib. 2021. Integrin-alpha(V)-mediated activation of TGF-beta regulates anti-tumour CD8 T cell immunity and response to PD-1 blockade. Nat Commun 12:5209.

30. McWilliam, H.E., S.B. Eckle, A. Theodossis, L. Liu, Z. Chen, J.M. Wubben, D.P. Fairlie, R.A. Strugnell, J.D. Mintern, J. McCluskey, J. Rossjohn, and J.A. Villadangos. 2016. The intracellular pathway for the presentation of vitamin B-related antigens by the antigen-presenting molecule MR1. Nat Immunol 17:531–537.

31. Meermeier, E.W., B.F. Laugel, A.K. Sewell, A.J. Corbett, J. Rossjohn, J. McCluskey, M.J. Harriff, T. Franks, M.C. Gold, and D.M. Lewinsohn. 2016. Human TRAV1-2-negative MR1-restricted T cells detect S. pyogenes and alternatives to MAIT riboflavin-based antigens. Nat Commun 7:12506.

32. Meng, K., J. Song, F. Qi, J. Li, Z. Fang, and L. Song. 2024. MT1G promotes iron autophagy and inhibits the function of gastric cancer cell lines by intervening in GPX4/SQSTM1. Sci Rep 14:28539.

33. Pampliega, O., I. Orhon, B. Patel, S. Sridhar, A. Diaz-Carretero, I. Beau, P. Codogno, B.H. Satir, P. Satir, and A.M. Cuervo. 2013. Functional interaction between autophagy and ciliogenesis. Nature 502:194–200.

34. Qu, J., B. Wu, L. Chen, Z. Wen, L. Fang, J. Zheng, Q. Shen, J. Heng, J. Zhou, and J. Zhou. 2024. CXCR6-positive circulating mucosal-associated invariant T cells can identify patients with non-small cell lung cancer responding to anti-PD-1 immunotherapy. J Exp Clin Cancer Res 43:134.

35. Reantragoon, R., A.J. Corbett, I.G. Sakala, N.A. Gherardin, J.B. Furness, Z. Chen, S.B. Eckle, A.P. Uldrich, R.W. Birkinshaw, O. Patel, L. Kostenko, B. Meehan, K. Kedzierska, L. Liu, D.P. Fairlie, T.H. Hansen, D.I. Godfrey, J. Rossjohn, J. McCluskey, and L. Kjer-Nielsen. 2013. Antigen-loaded MR1 tetramers define T cell receptor heterogeneity in mucosal-associated invariant T cells. J Exp Med 210:2305–2320.

36. Ruf, B., M. Bruhns, S. Babaei, N. Kedei, L. Ma, M. Revsine, M.R. Benmebarek, C. Ma, B. Heinrich, V. Subramanyam, J. Qi, S. Wabitsch, B.L. Green, K.C. Bauer, Y. Myojin, L.T. Greten, J.D. McCallen, P. Huang, R. Trehan, X. Wang, A. Nur, D.Q. Murphy Soika, M. Pouzolles, C.N. Evans, R. Chari, D.E. Kleiner, W. Telford, K. Dadkhah, A. Ruchinskas, M.K. Stovroff, J. Kang, K. Oza, M. Ruchirawat, A. Kroemer, X.W. Wang, M. Claassen, F. Korangy, and T.F. Greten. 2023. Tumor-associated macrophages trigger MAIT cell dysfunction at the HCC invasive margin. Cell 186:3686–3705 e3632.

37. Shi, L., J. Lu, D. Zhong, M. Song, J. Liu, W. You, W.H. Li, L. Lin, D. Shi, and Y. Chen. 2023. Clinicopathological and predictive value of MAIT cells in non-small cell lung cancer for immunotherapy. J Immunother Cancer 11:

38. Stepp, M.W., R.J. Folz, J. Yu, and I.N. Zelko. 2014. The c10orf10 gene product is a new link between oxidative stress and autophagy. Biochim Biophys Acta 1843:1076–1088.

39. Vacchini, A., A. Chancellor, Q. Yang, R. Colombo, J. Spagnuolo, G. Berloffa, D. Joss, O. Oyas, C. Lecchi, G. De Simone, A. Beshirova, V. Nosi, J.P. Loureiro, A. Morabito, C. De Gregorio, M. Pfeffer, V. Schaefer, G. Prota, A. Zippelius, J. Stelling, D. Haussinger, L. Brunelli, P. Villalta, M. Lepore, E. Davoli, S. Balbo, L. Mori, and G. De Libero. 2024. Nucleobase adducts bind MR1 and stimulate MR1-restricted T cells. Sci Immunol 9:eadn0126.

40. VandeKopple, M.J., J. Wu, E.N. Auer, A.J. Giaccia, N.C. Denko, and I. Papandreou. 2019. HILPDA Regulates Lipid Metabolism, Lipid Droplet Abundance, and Response to Microenvironmental Stress in Solid Tumors. Mol Cancer Res 17:2089–2101.

41. Wong, E.B., T. Ndung’u, and V.O. Kasprowicz. 2017. The role of mucosal-associated invariant T cells in infectious diseases. Immunology 150:45–54.

42. Zou, J., and N. Shankar. 2016. The opportunistic pathogen Enterococcus faecalis resists phagosome acidification and autophagy to promote intracellular survival in macrophages. Cell Microbiol 18:831–843.

