## supplemental figures for "Synergistic role of riboflavin-auxotrophic *Enterococcus* for MR1 expression and intra-tumoral mucosal associated invariant T (MAIT) cell activation"

**Supplementary information**

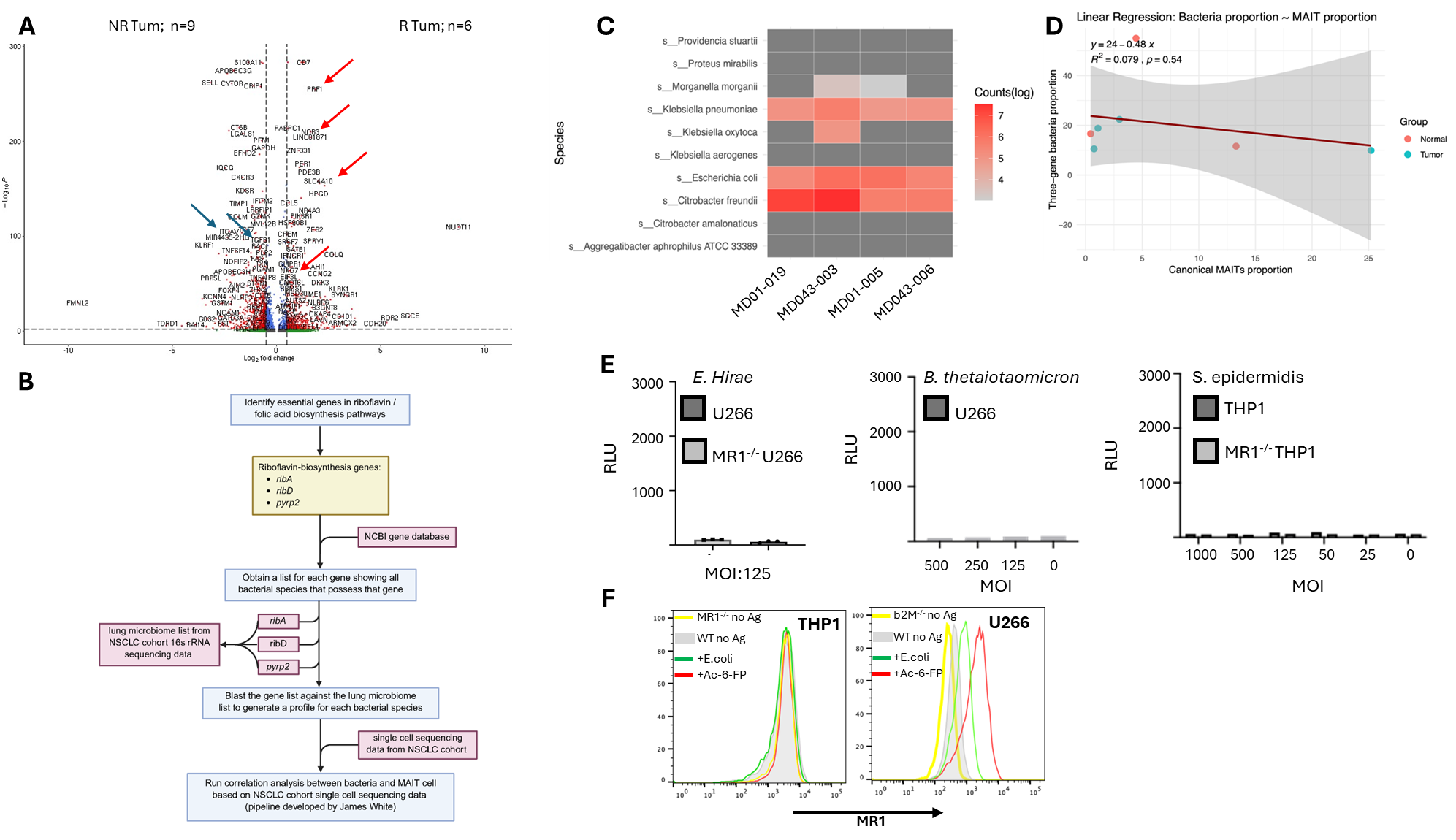

**Supplementary figure S1. A.** Volcano plot depicting differentially expressed genes in *cytotoxic MAIT 1* cluster in tumor tissues from responder (n = 6) vs Non-responder (n = 9) patients. **B,** Schematic representation of the pipeline predicting the production of the 5-A_RU by the intratumoral microbiota. **C**, heat map representing the abundance of 5-A-RU producing bacteria among all bacteria in lung tumor samples. **D**, linear regression analysis showing the absence of correlation between the proportion of intratumoral 5-A-RU-producing bacteria and proportion of intratumoral conventional MAIT (TCRAV1-2/J33,20,12). **E**, Riboflavin auxotrophic *E. hirae*, riboflavin-producing *B. thetaiotaomicron* and *S. epidermis* do not activate MAIT TCR MD01-005 J33-4. **F**, flow cytometry analysis of MR1 expression on THP1 (Left) and U266 (Right) without antigen (grey) or in presence of *E. coli* (green), *Ac-6-FP* (red). MR1 negative MR1^-/-^ THP1 and β2m^-/-^ U266 (yellow) are used as negative control.

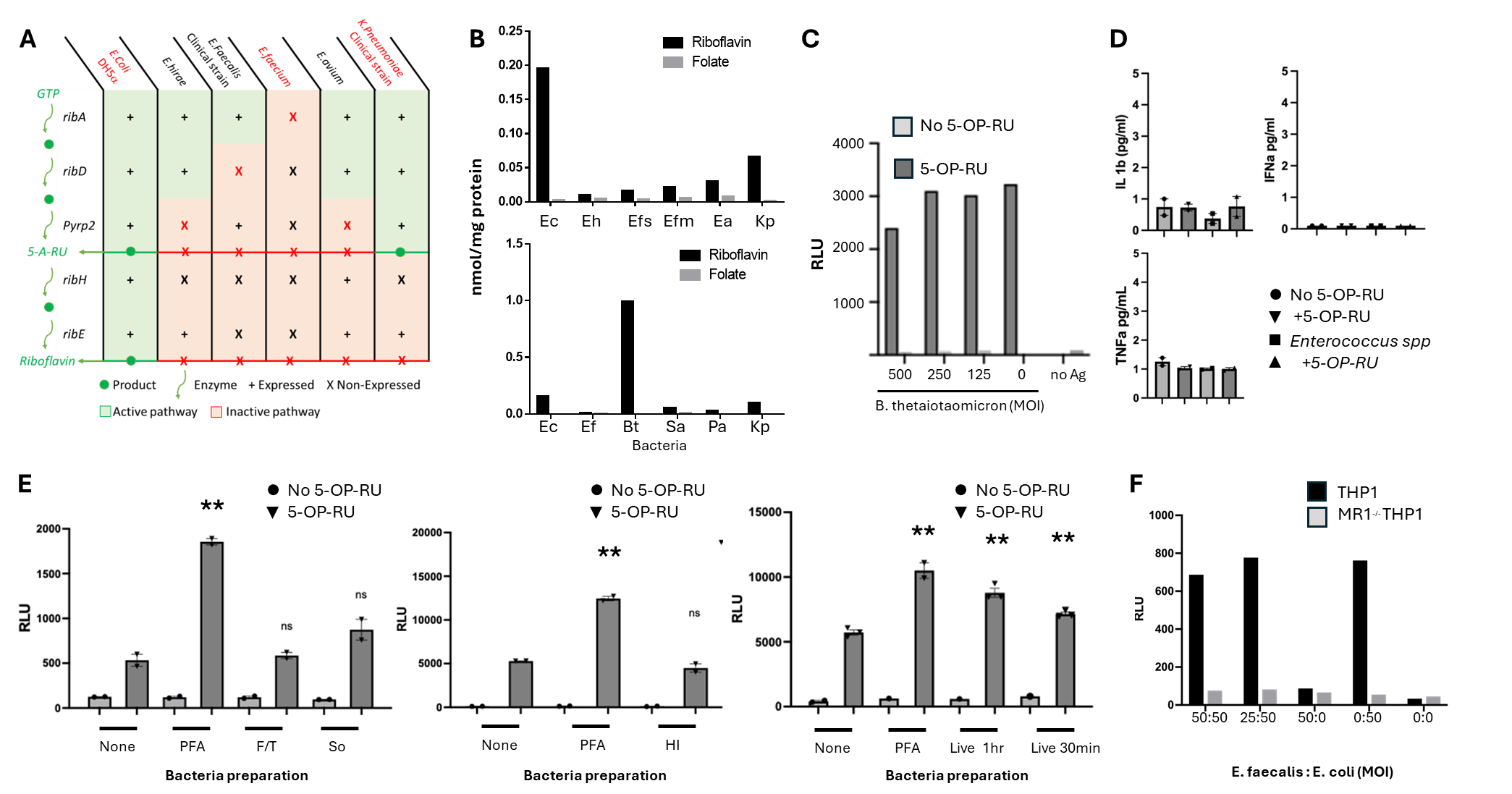

**Supplementary figure S2. A**, Bacteria strains used in our tests were sequenced to determine the expression of genes encoding the enzymes responsible for the biosynthesis of Riboflavin from GTP. +, presence of the gene; X, absence of the gene. Red boxes, deficient metabolic steps; Green, proficient metabolic steps. **B**, metabolomic analysis of the riboflavin biosynthesis by *E.faecalis* (EF), *E.coli* (EC), *Staphylococcus aureus* (SA), *Pseudomonas aeruginosa* (PA) to validate the genomic analysis in **A**. **C**, MAIT TCR J33-4 recognition of U266 cells incubated overnight with (dark grey) or without (light grey) 5-OP-RU, in presence or not of decreasing MOI of *B. thetaiotaomicron*. TCR activation is measured by bioluminescence reading. **D**, INFα, IL1β and TNF measurements by ELISA, in the culture supernatants of U266 cultured (inversed black triangle) or not (black circle) with 5-OP-RU, in presence of *Enterococcus* (black triangle) or not (black square). **E**, MAIT TCR J33-4 recognition of U266 cells incubated overnight with (inverse black triangles) or without (black circles) 5-OP-RU, in presence or not of E. coli, PFA fixed (PFA), heat inactivated (HI), sonicated (So), Frozen/Thawed lysed (F/T) or alive. **F**, absence of synergy by *E. faecalis* for TCR J33-1 activation by THP1 cells cultured in the presence of PFA fixed *E. coli* at different MOIs. MR1^-/-^ THP1 cells were used as negative control. All data are shown as mean ± SEM with technical replicates represented by number of symbols in each graph. ****:P<0.0001, ***:0.0001<P<0.001, **:0.001<P<0.01, *:0.01<P<0.05 , ns (non significant) by unpaired t-test (two-tailed).

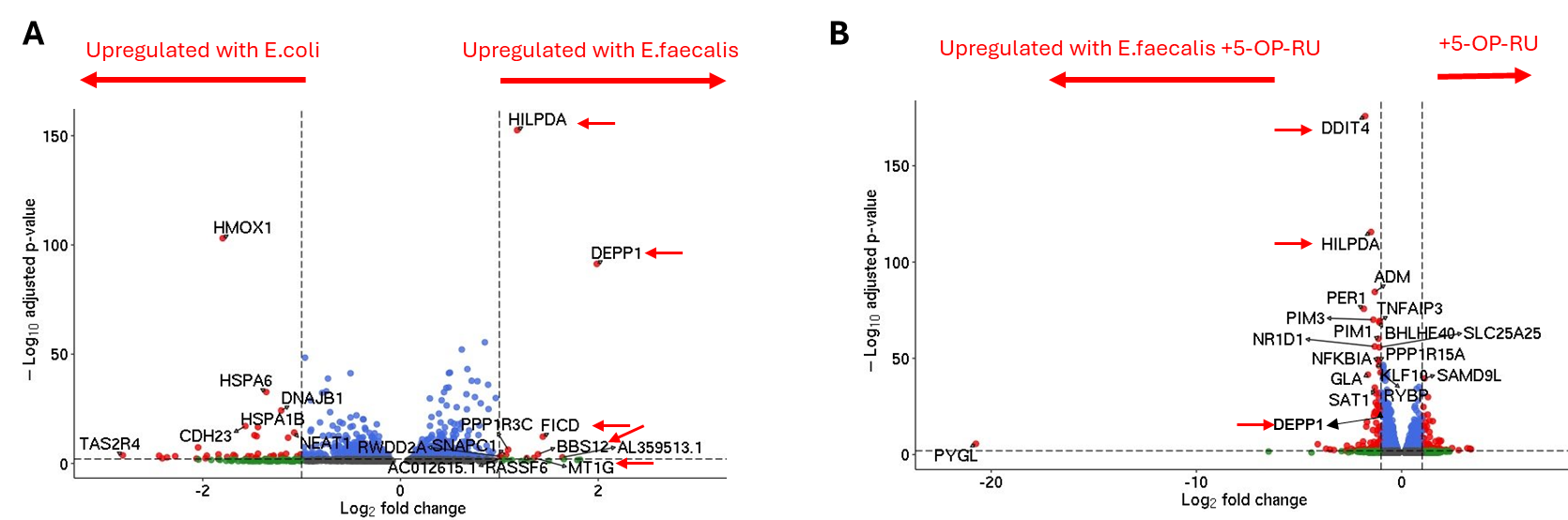

**Supplementary figure S3. A&B,** Volcano plots representing RNAseq analysis of U266 cell line incubated with *E. faecalis* compared to E. coli (**A**) and 5-OP-RU + *E.* faecalis compared to 5-OP-RU alone (**B**). Significant genes were selected when p<0.01 and -Log2(FC>2). Red arrows indicate genes associated with autophagy.
